# Gone but not forgotten: dynamics of sperm storage and potential ejaculate digestion in the black soldier fly *Hermetia illucens*

**DOI:** 10.1101/2024.05.31.596011

**Authors:** Frédéric Manas, Harmony Piterois, Carole Labrousse, Laureen Beaugard, Rustem Uzbekov, Christophe Bressac

**Affiliations:** Institut de Recherche sur la Biologie de l’Insecte (IRBI), UMR CNRS 7261 Université de Tours, 37200 Tours, France; Plateforme IBiSA de Microscopie Electronique, Université de Tours et CHRU de Tours, 37200 Tours, France; Faculty of Bioengineering and Bioinformatics, Moscow State University, 119991, Moscow, Russia

**Keywords:** spermathecae, sperm viability, successive egg-laying, sperm plug, nuptial gift

## Abstract

When it comes to describe reproduction in internally fertilized species, understanding the dynamics of sperm storage is crucial to unravel the complexity of post-copulatory sexual selection processes. This physiological process goes from sperm transfer to its use for fertilization. In this framework, the spatiotemporal dynamics of sperm storage were described in the black soldier fly (BSF) with fluorescence and transmission electron microscopy (TEM). Besides being of great economic interest, BSF females have highly compartmentalized spermathecae. Spermatozoa were counted both during and after mating in two successive spermathecae compartments: the fishnet canals and the reservoirs. In addition to seminal fluids, male transfers a sperm plug in the fishnet canals, then only some spermatozoa reach the reservoirs over 2 days. TEM observations of the fishnet canals revealed potential digestive functions, explaining the decline in the number and the viability of spermatozoa that have not been stored in the reservoirs. After one mating, females laid up to three fertile clutches, evidencing no constraints on sperm quantity or quality. Spermatic and ultrastructural investigations strongly suggest that BSF ejaculate acts both as a sperm plug and a as nuptial gift, reinforcing the interest in studying this insect as a model for sexual selection.

## 1. Introduction

In species with internal fertilization, sperm storage—from sperm transfer to its use in fertilization—is a fundamental process of sexual reproduction, as it is at the core of post-copulatory selection processes, i.e.,sperm competition [1] and cryptic female choice [2]. As a result, the spatiotemporal dynamics of sperm storage are of great importance to understand sexual selection [3–5]. The process of sperm storage includes three successive steps, involving both the male and the female [6]: 1. Sperm transfer from the male reproductive tract to the receptive compartment of the female. 2. Migration of sperm to the site of storage or future fertilization. 3. Use of only some spermatozoa for fertilization. At each step, spermatozoa could be selected by mechanisms involving sperm competition or/and cryptic female choice [7]. Thus in species where females have sperm storage organs, such as in most arthropods [8], dynamics of sperm storage and use can be complex. In this respect, the study of both temporal and spatial [3,4] dynamics of sperm storage allows to highlight the processes through which post-copulatory sexual selection occurs.

Sperm precedence is one of these selective processes—whether it is first male or last male sperm precedence—which is the way through which a male sperm is favored for fertilization according to the relative order of arrival in the female’s reproductive organs [9]. First male sperm precedence is often due to the transfer of “mating plugs” formed by spermatozoa [10], coagulation of seminal fluids, or male genitalia [11–13]. In contrast, last male sperm precedence, can result from mechanisms of sperm displacement [3] or sperm stratification [14] favoring the last copulating male. The role of the female is obviously not to be overlooked here as sperm dumping/ejection is a way to favor paternity of some males over others—i.e., to perform cryptic female choice [2,15].

After mating and storing spermatozoa, the female will use them to fertilise oocytes. Consequently, there is a decrease in the amount of stored spermatozoa, which can lead to sperm limitation in some species—i.e. spermatozoa being a limited resource for the female [16]. This constraint can rise from both quantity and quality, as spermatozoa may degrade over time in storage [17]. Sperm limitation not only affects egg production [18] but is also intricately tied to sexual selection as responses to sperm limitation may include selection for males producing/transferring more sperm, as well as females remating behaviour [19–21].

Among insects, the black soldier fly (BSF, *Hermetia illucens*, Diptera, Stratiomyidae) is the source of a lot of economic interests [22,23]. However, the understanding of the biology of this species, particularly in its adult stage, remains limited [24]. It has been shown that BSF exhibits polygynandrous mating system [25], are capable of laying multiple clutches [25] and that males respond to the risk of sperm competition by adjusting their production and allocation of spermatozoa depending on social contexts [26]. During their immobile mating, adults are attached from 20 minutes to more than one hour [27]. BSF shows an interesting reproductive tract [28]. In females, each of the thin bases of the three separated spermathecae enlarge in a longer canal with a musculated outer wall and a fishnet-ornemented inner matrix (figure 1A, 1B)[28]. Then, a thinner canal embedded with muscles leads to a thin rigid rod, making an elbow with a second rigid rod pierced with small holes preceeding the sperm reservoir. Obviously, such a complex structure raises questions about the reproductive processes of BSF. Recently, it has been demonstrated that mating in BSF enhances female longevity [29], implying the possible transfer of nuptial gifts. However, it is noteworthy that BSF does not transfer spermatophylaxes or spermatophores during mating [28]. Therefore, if males do indeed provide a nuptial gift, it has to be investigated in the whole ejaculate—spermatozoa and seminal fluids—as it would likely be in the form of an endogenous genital gift [30], potentially including non-nutritional products akin to those found in *Drosophila melanogaster* seminal fluids [31].

**Figure 1.**
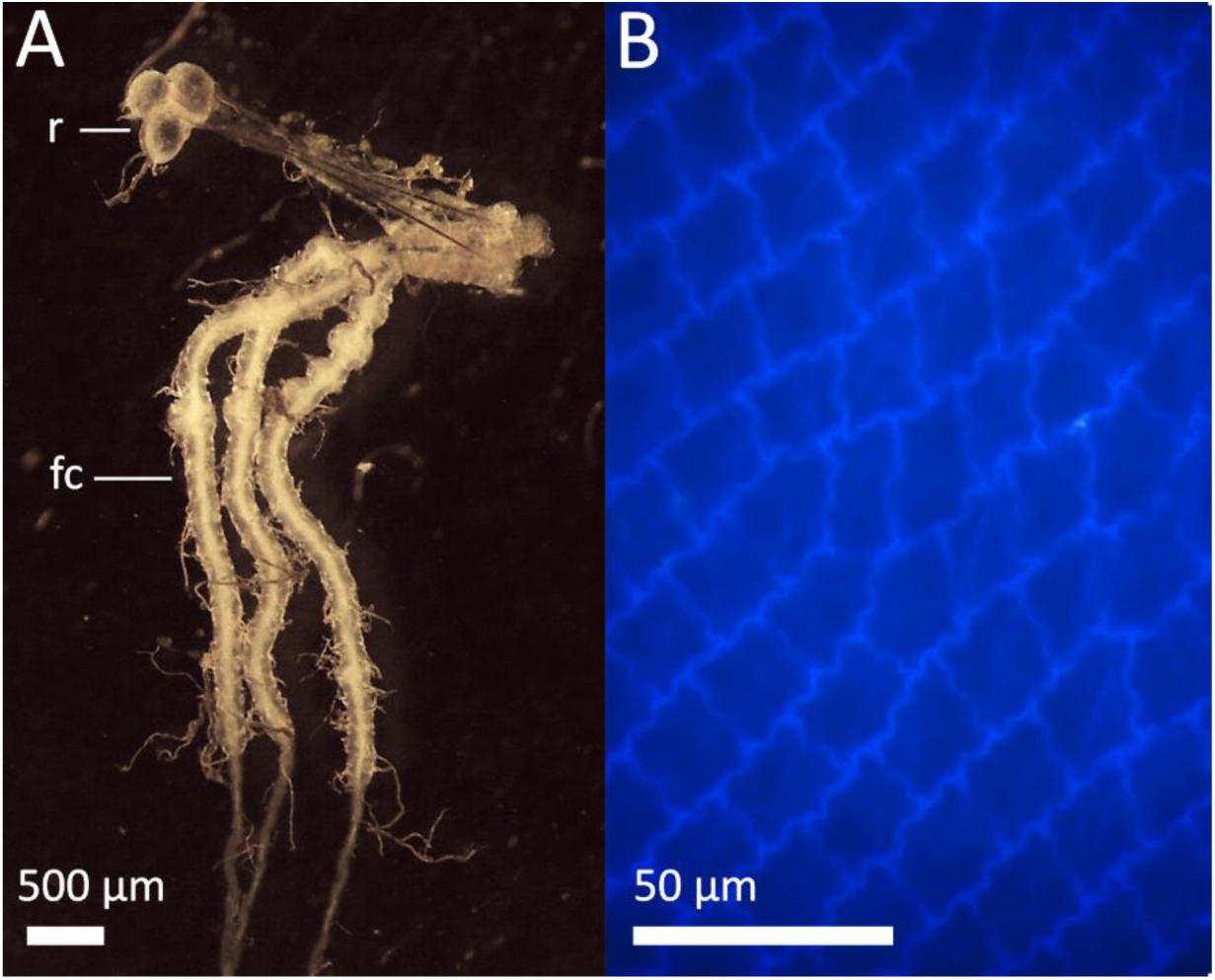
(A) The three spermathecae of a virgin female BSF and (B) the fishnet canal matrix under a fluorescence microscope. The color of the matrix in (B) is due to autofluorescence under UV light. Abbreviations: *r* reservoirs, *fc* fishnet canals.

Using a lab population of BSF, the present study aimed to investigate: (1) the spatiotemporal dynamics of sperm storage – from transfer to use; (2) the variations in sperm quality along its storage in females; (3) the ultrastructure of the spermathecae in relation to the mating status of the female; and (4) the possibility of a nuptial gift transfer during mating.

## 2. Materials and methods

### (a) Rearing conditions, mating and size measures

BSF were reared and isolated as described in [28]. As BSF will not initiate mating when a single pair is placed in a cage (personal observations), first step to get matings consisted in transferring 20 virgin males to 15×15×15 cm cages containing 20 virgin females. Individuals remained in contact for 3 hours to obtain mating pairs. Different series of flies were used for sperm storage dynamics description (15 series), the relationship between male size and fishnet canals volume (6 series), clutches monitoring (13 series), viability assessments (6 series) and transmission electron microscopy (1 serie).

To measure their head width, BSF were photographed under a Nikon SMZ745T stereomicroscope (x3.35 magnification) (Nikon, Japan) with a Leica IC 80 HD camera (Leica, Germany) using ImageJ. Head width can be considered as a reliable proxy of the size of the individuals [32].

### (b) Sperm storage dynamics

To describe the dynamics of sperm storage before females began laying eggs, flies were dissected (n= 128) at different intervals during and after mating. Preliminary observations showed that mating lasted mean ± SE = 33.32 ± 10.19 minutes (n = 166) in the experimental rearing that has been used for these experiments. Considering this mating duration, females were dissected 5 (n = 5), 10 (n = 15), 15 (n = 15), 20 (n = 15), 25 (n = 14), 30 (n = 15) minutes after the beginning of mating and just after the end of mating (n = 12). Other females were dissected 24 hours (n = 15) and 48 hours (n = 10) after mating, before oviposition, and after their first egg-laying (n = 12). Petri dishes in which females were isolated were examined with DAPI under a fluorescence microscope to search for potential dumped sperm. Some females (n = 8) were also kept in “cages” made of microscope slides (as in [33]) to search for dumped sperm.

Another set of females (n = 47) was dissected at the end of mating to describe the relationship between male size and the volume of the spermathecae just after mating.

For all females, the abdomen was opened to collect the three spermathecae which were then placed on a microscope slide in a drop of PBS (Phosphate Buffer Saline). After dissections, spermathecae were photographed to take measures of the fishnet canals width and length with imageJ in order to estimate their volume using the formula of a cylinder: *V = π.r².h*. First measures showed that volumes of the three fishnet canals were higly similar within a female (using the coefficient of variation (CV) which is calculated as: CV = SD/mean x 100, mean CV = 8.67 ± 5.20, n = 13) so at least one fishnet canal was measured and individuals for which one or two spermathecae were lost during dissection were still considered.

After taking a picture, a drop of DAPI was applied, then the spermathecae were crushed with a microscope coverslip to release spermatozoa and DAPI-labelled nuclei of spermatozoa were counted under a fluorescence microscope (Olympus CX40, Japan) with a x20 objective. In order to describe the spatial distribution of spermatozoa, the spermathecae were divided in 3 areas in which spermatozoa were counted: the first half of the fishnet canals, the second half of the fishnet canals and the reservoirs (figure 1A). Spermatozoa counts were realized by visual analysis as in [26,28] and were repeatable (Pearson correlation coefficients: r = 0.91, P < 0.001, n = 12 samples counted two times). For some females dissected just at the end of mating (n = 13), spermatozoa were counted in the three spermathecae to ensure the number of spermatozoa transferred to each spermathecae was not variable (see results). Then, the number spermatozoa were counted in one spermatheca and multiplied by 3 to estimate the total number of spermatozoa transferred. Sperm stored two days after mating and after the first egg-laying were counted in the three reservoirs.

### (c) Clutches monitoring

To evaluate how many eggs a female could lay, mating pairs (n = 114) were gently placed on the lid of a petri dish and taken out of the cage until the end of mating. Time was recorded once matings were completed to measure duration, and pairs were kept together at 24°C with a cotton saturated with water in petri dishes where extra mating are not possible [28]. No food was provided to ensure that all females were on the same conditions for egg laying and to prevent mold.

Pairs were observed every day to check for eggs. To collect eggs, females and males were transferred to a new petri dish with a new cotton soaked with water. The eggs were gently separated with forceps to be counted by visual analysis under a Nikon SMZ745T stereomicroscope (x3.35 magnification) (Nikon, Japan) and then put in an incubator to check the fertility of the clutches at 27°C which was assessed by the visualization of the embryo red eye-spots before eclosion (figure S1). Fertility was assessed before hatching because larvae that just eclosed disperse very quickly, which induces a mixture of eggs that is difficult to detect as hatched or not hatched. This process was repeated until the death of the female which was then dissected. For this clutches monitoring, spermatozoa were not counted; the reservoirs were examined to determine if spermatozoa were still present and to estimate the order of magnitude of the number of remaining spermatozoa.

### (d) Sperm viability assessment

Sperm viability in the fishnet canals was measured at two stages to assess if it could decrease after mating, before reaching the reservoirs. The assessment was conducted directly after mating (n = 25) and one day after mating (n = 26). Similarly, to evaluate whether sperm quality could be a factor limiting egg-laying, sperm viability in female reservoirs was measured at three stages: 2 days (n = 18), 7 days (n = 10), and 14 days (n = 9) after mating.

Viability of spermatozoa was assessed using a Live/Dead Sperm Viability Kit (Invitrogen). After dissection of the female, spermathecae were opened with forceps on a glass slide to be dipped in 5 μL of a 1: 20 dilution of 1 mM SYBR-14. After 10 minutes of incubation in the dark, 5 μL of propidium iodide were added to the preparation. Pictures of the three fishnet canals/reservoirs were taken under a fluorescence microscope to blind count green and red sperm nuclei. Spermatozoa that were half-green half-red were counted as dead (figure S2).

### (e) Transmission electron microscopy

Spermathecae from females of different mating statuses were investigated for ultrastructure through TEM: a virgin female, a female that mated on the day of its dissection, and a female that mated two days before its dissection. For each of them, semi-thin sections of their spermathecae—fishnet canals and reservoirs—were realized to describe their structure.

Spermathecae samples were fixed in mixture of 2% paraformaldehyde (Merck, Darmstadt, Germany), 2% glutaraldehyde (Agar Scientific, France) and 0.1 M sucrose in 0.1 M cacodylate buffer (pH 7.4) for 24 hours, washed three × 30 min in 0.1 M of cacodylate buffer, and post-fixed for 1.5 hours with 2% osmium tetroxide (Electron Microscopy Science, USA) in 0.1 M cacodylate buffer. After washing in 0.1 M cacodylate buffer for 20 min and two x 20 min in distillated H2O, samples were dehydrated in a graded series of ethanol solutions (50% ethanol two x 15 minutes; 70% ethanol two x 15 minutes and third portion of 70% ethanol for 14 hours; 90% ethanol two x 20 minutes; and 100% ethanol three x 20 minutes). Final dehydration was performed by 100% propylene oxide (PrOx, VWR Int., France) three x 20 min. Then, samples were incubated in PrOx/EPON epoxy resin (Fluka, Switzerland) mixture in a 2:1 ratio for two hours with closed caps, in PrOx/EPON epoxy resin (Fluka, Switzerland) mixture in a 1:2 ratio for two hours with closed caps and 1.5 hours with open caps, and in 100% EPON for 16 h at room temperature. Samples were replaced in new 100% EPON and incubated at 37 °C for 24 hours and at 60 °C for 48 hours for polymerization.

Semi-thin sections (thickness 0.8 µm) were cut with a “Leica Ultracut UCT” ultramicrotome (Leica Microsysteme GmbH, Wien, Austria), placed on glass, stained with Toluidine blue (Electron Microscopy Science, Hatfield, PA, USA) and embedded in Epon resin (Fluka, Switzerland) which was allowed to polymerize for 48 hours at 60°C. The sections were then observed with Nikon Eclipse 80i microscope (Nikon, Japan) connected to DS-Vi1 camera driven by Nis-Element D 4.4 imaging software (Nikon, Japan).

Ultra-thin sections (thickness 70 nm) were placed on TEM one-slot grids (Agar Scientific, France) coated with Formvar film and stained 20 minutes with 5% uranyl acetate (Electron Microscopy Science, Hatfield, PA, USA) and 5 minutes Reynolds lead citrate. The sections were then observed at 100 kV with a Jeol 1011 TEM (Tokyo, Japan) connected to a Gatan digital camera driven by Digital Micrograph software (GMS 3, Gatan, Pleasanton, CA, USA).

### (f) Statistical analyses

Wilcoxon-Mann-Whitney tests were used to compare the volume of fishnet canals and the number of spermatozoa counted during mating, as the data were not parametric. A Benjamini-Hochberg correction was applied to adjust p-values [34,35] due to multiple comparisons across different dissection timings.

To assess variability in the number of laid eggs, a linear regression was performed with the log-transformed total number of eggs laid as the dependent variable. Covariates included male and female head width, the number of clutches of the female, and mating duration.

Another linear regression was performed to analyze the mean volume of the fishnet canals just after mating. Covariates in this model included male and female head width, mating duration, and the number of spermatozoa transferred.

A third linear regression was carried out to study female longevity. Covariates in this model were male and female head width, male longevity, and mating duration.

A Wilcoxon-Mann-Whitney test was used to compare sperm viability percentages in the fishnet canals directly after mating and one day after. A Kruskal-Wallis test was used to compare sperm viability percentages in the reservoirs among females dissected 2 days, 7 days, and 14 days after mating. A paired-sample t-test was used to compare the number of eggs between first and second clutches.

All statistical analyses were performed using R version 4.0.2 (R Core Team, 2020). The significance level was set at alpha = 0.05 for all tests. Residual uniformity of the linear models was assessed using the performance package [36]. Quantitative data were presented as mean ± standard deviations (SD) when they were normally distributed and Q25 (first quartile); median; Q75 (third quartile) when they were not.

## 3. Results

### (a) Sperm storage dynamics

At the early times of mating – 5 to 10 minutes after the beginning, the first part of the female spermathecae—the fishnet canals—were empty of seminal fluid (1.88 ± 0.86 μL, figure 2A) and almost empty of spermatozoa (Q25 = 0; median = 0; Q75 = 0 spermatozoa in the fishnet canals with n = 14 females having no spermatozoa and n= 1 female having 8 spermatozoa (figure 3A)). After 15 minutes of mating, fishnet canals were swelled by seminal fluids transferred by the male (4.34 ± 2.02 μL, figure 2A) and reached their maximum volume between 25 minutes and the end of mating (7.79 ± 1.94 μL, figure 2A, S3). When taken out from the inside of the female tract, the handling of this seminal fluid with forceps made it looks more like a jelly than a fluid (figure S4).

**Figure 2.**
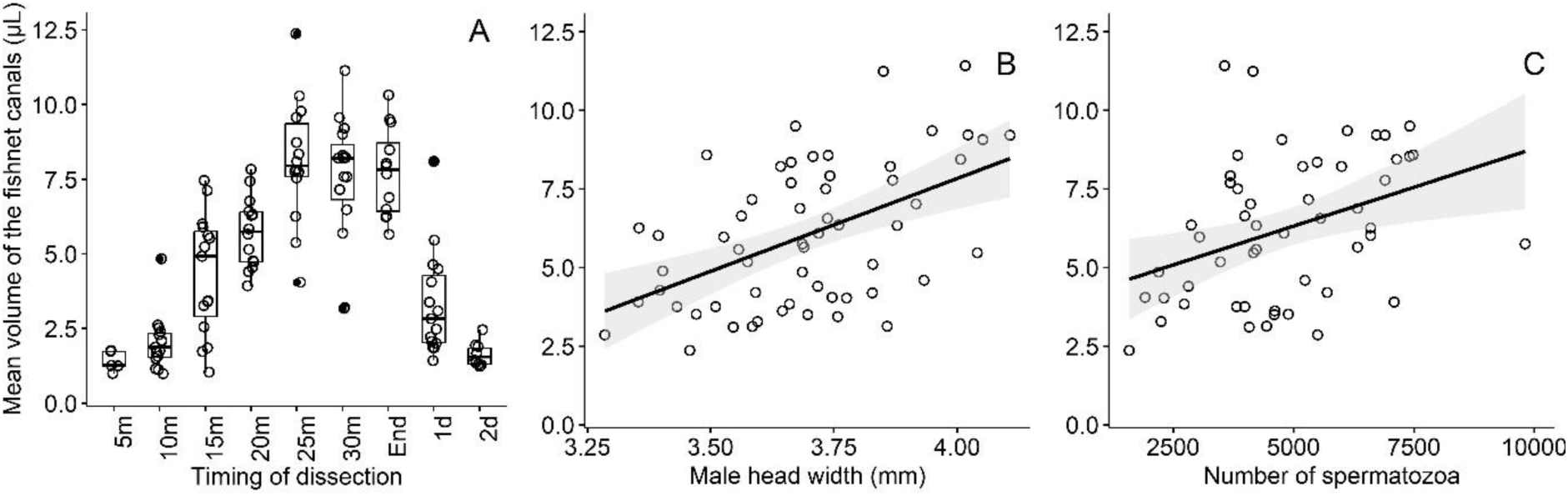
Mean fishnet canals volume in μL according to (A) the timing of dissection and to (B) male head width when measured at the end of mating and (C) the number of spermatozoa in the fishnet canals at the end of mating. For (A), box plots show median (horizontal bars), upper, and lower quartiles (borders of the box). Whiskers extend from the 10th to the 90th percentiles. For (B) and (C) lines represent linear regression and shaded areas represent confidence intervals.

**Figure 3.**
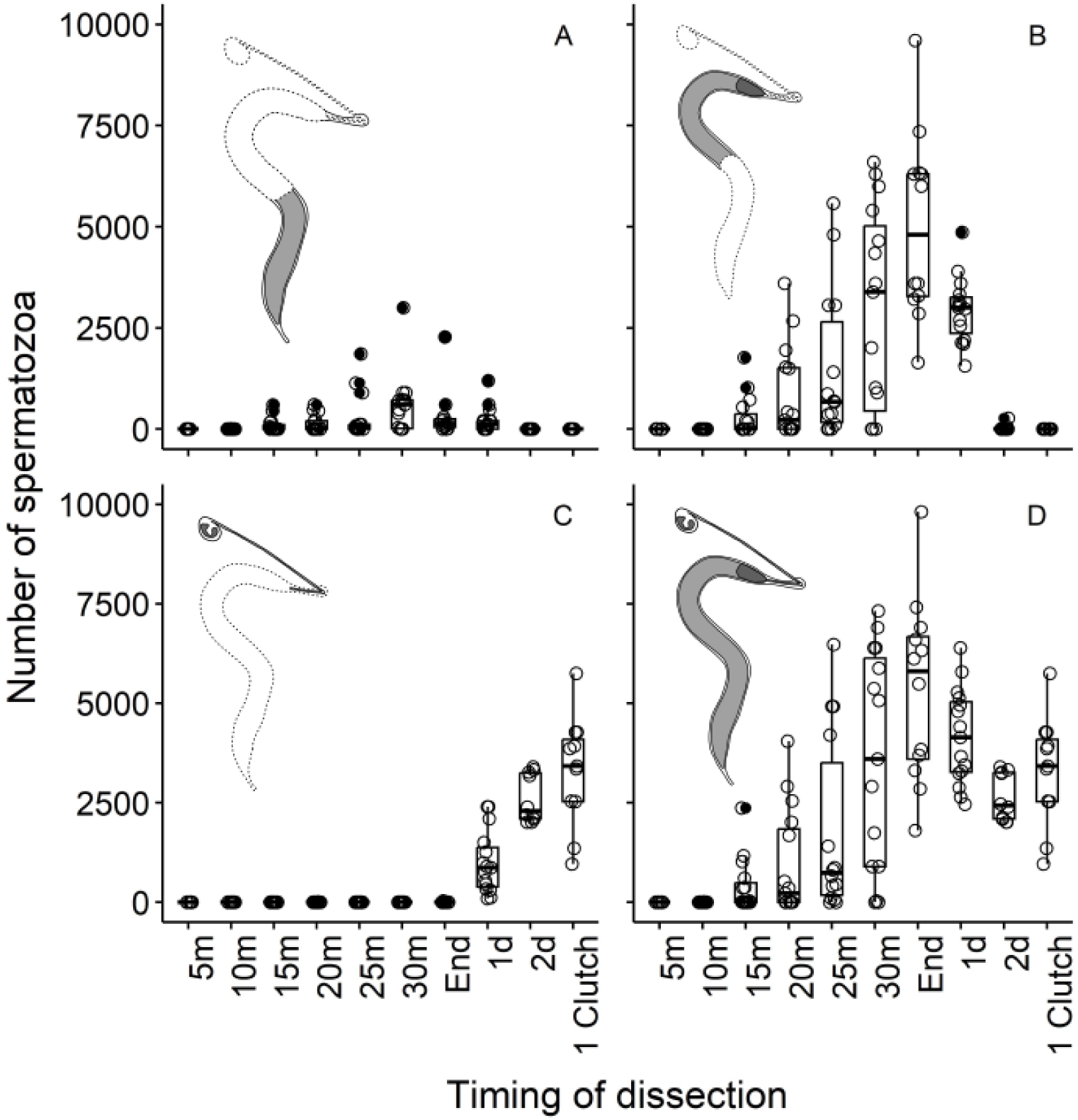
Number of spermatozoa according to the timing of dissection considering the three spermathecae (A) in the first half of the fishnet canals, (B) in the second half of the fishnet canals, (C) in the reservoirs of the spermathecae and (D) in the totality of the spermathecae. Box plots show median (horizontal bars), upper, and lower quartiles (borders of the box). Whiskers extend from the 10th to the 90th percentiles.

When measured at the end of mating, the mean volume of the fishnet canals was significantly and positively correlated to male head width (linear regression: n = 47, F_1,42_ = 26.70, P < 0.001, full model R² = 0.43, β ± SE = 6.03 ± 1.43, Fig.2B) and to the number of spermatozoa transferred (F_1,42_ = 6.67, P = 0.01, β ± SE = 4.04 ± 1.56, figure 2C). However, no significant relationship was found between the mean volume of the fishnet canals and female head width (F_1,42_ =3.00, P = 0.09) or mating duration (F_1,42_ = 2.26, P = 0.14).

The number of spermatozoa transferred to each spermathecae (1659.5 ± 548.37 spermatozoa) was relatively similar acrross the three fishnet canals of a female (mean CV for the number of spermatozoa in spermathecae of a female = 15.36 ± 8.17)(figure S5).

Spermatozoa were visible in the fishnet canals from some minutes upwards (at 15 minutes: 395 ± 635.18, Q25 = 0; median = 5; Q75 = 485 spermatozoa in the fishnet canals, figure 3A, 3B, 3D) and their number increased until the end of mating to reach a mean of 5346 ± 2290 (figure 3D). Most of the spermatozoa accumulated at the top of the fishnet canals (figure 3C, 3D) – 336.2 ± 635.2, Q25 = 0; median = 135; Q75 = 255 spermatozoa in the first half of the fishnet canals and 5007.5 ± 2309.857 spermatozoa in the second half of the fishnet canals at the end of mating (Wilcoxon test: P < 0.001) - and no spermatozoa were observed in the reservoirs until the day after mating (figure 2B).

The total number of spermatozoa began to decrease 24 hours after mating (4168 ± 1198.2 spermatozoa considering the entire spermathecae) and became significantly lower (Wilcoxon test: P < 0.01) 48 hours after the end of mating even though females had not laid (2634 ± 601.7, Q25 = 2100; median = 2430; Q75 = 3262 spermatozoa). However, no dumped sperm was found in the petri dishes or in microscope slides cages containing females. 24 hours after mating, the spatial distribution of spermatozoa changed and the reservoirs began storing spermatozoa (spermatozoa in the reservoirs at the end of mating: 2.50 ± 8.66, Q25 = 0; median = 0; Q75 = 0 and 24 hours after mating: 1000 ±783.4; Wilcoxon test: P < 0.001, figure 3). In the same time, fishnet canals became progressively empty from spermatozoa (spermatozoa in the second half of the fishnet canals at the end of mating: 5007.5 ± 2309.86 and 24 hours later: 2948 ± 816.44, Wilcoxon test: P = 0.01, figure 3B) and seminal fluid (mean volume at the end of mating: 7.74 ± 1.49 μL and 24 hours later: 3.33 ± 1.78 μL, Wilcoxon test: P < 0.001) (figure 2A).

After a first egg-laying, the number of spermatozoa was not significantly different between the three reservoirs (Anova: Df = 1, F = 0.08, P = 0.78), but the coefficient of variation of the number of spermatozoa across the reservoirs of a same female was high (mean cofficient of variation for the number of spermatozoa in the reservoirs of a female = 33.01 ± 27.63). No significant loss of spermatozoa has been observed after the first egg laying with 2634 ± 601.67, Q25 = 2100; median = 2430; Q75 = 3262 spermatozoa 48 hours after mating and 3041.2 ± 1578.7 spermatozoa after first egg-laying.

### (b) Clutches monitoring

The mean of the total number of eggs laid by females after mating was 866.4 ± 355.7. From 114 matings, 47 single, 62 double as well as 2 triple clutches were obtained and 3 females didn’t lay at all. The total number of eggs laid varied significantly and positively with female head width (linear regression: n = 108, F_1,103_ = 32.81, P < 0.001, full model R² = 0.47, β ± SE = 0.35 ± 0.18), and negatively with male head width (F_1,103_ = 9.83, P < 0.01, β ± SE = −0.49 ± 0.16). Mating duration was not linked to the total number of eggs laid (F_1,103_ = 0.18, P = 0.67).

Females laid their first clutches 2.88 ± 1.09 days after mating. They laid more eggs (paired student test: t = 5.35, P < 0.001) in their first clutch (569.6 ± 163.3) than they did in their second clutch (483.1 ± 156.5)(figure S6). Only two females laid third clutches containing respectively 85 and 505 eggs. On average, first clutches were laid at an age of 6.30 ± 1.93 days and second clutches at an age of 12.36 ± 2.44 days. The two third clutches were laid when the females had 12 and 11 days, which was 3 days after the second clutches for both of them. Their was no significant difference of fertility between first (89 % of fertile clutches) and second clutch (78 % of fertile clutches) (χ^2^ test: χ^2^ =1.49, Df = 1, P = 0.22). All the females still had spermatozoa in their reservoirs after laying their clutches.

Female longevity was significantly and positively related to her size (linear regression: n = 75, F_1,70_ = 9.80, P < 0.01, full model R² = 0.17, β ± SE = 3.89 ± 1.21). However, female longevity was not significantly related to the size of the male she mated with (F_1,70_ = 1.96, P = 0.16) nor with mating duration (F_1,70_ = 0.97, P = 0.33). There was a slight but significant (F_1,70_ = 6.80, P = 0.01) positive relationship between the longevity of the female and the longevity of the male she mated with (β ± SE = 0.14 ± 0.05).

### (c) Viability of the spermatozoa

In the fishnet canals, the proportion of viable spermatozoa was higher when counted directly at the end of mating compared to when counted one day after mating (Wilcoxon test: W = 646, P < 0.001), decreasing from 84 ± 13 % to 20 ± 17 % (figure 4A). In the reservoirs, the proportion of viable spermatozoa was high with a mean proportion of 90 ± 8 % at the three time intervals. The proportion of viable spermatozoa in the reservoirs was not significantly different (Wilcoxon test: P = 0.796) when counted 2 days after mating (89 ± 7 %) compared to when counted 7 days after mating (89 ± 9 %). In the same way it was not significantly different (Wilcoxon test: P = 0.14) between counts realized 7 days after matings and counts realized 14 days after mating (95 ± 5 %). However the proportion of viable spermatozoa counted 14 days after mating (95 ± 5 %) was significantly higher (Wilcoxon test: P = 0.03) than the one counted 2 days after mating (89 ± 7 %) (figure 4B).

**Figure 4.**
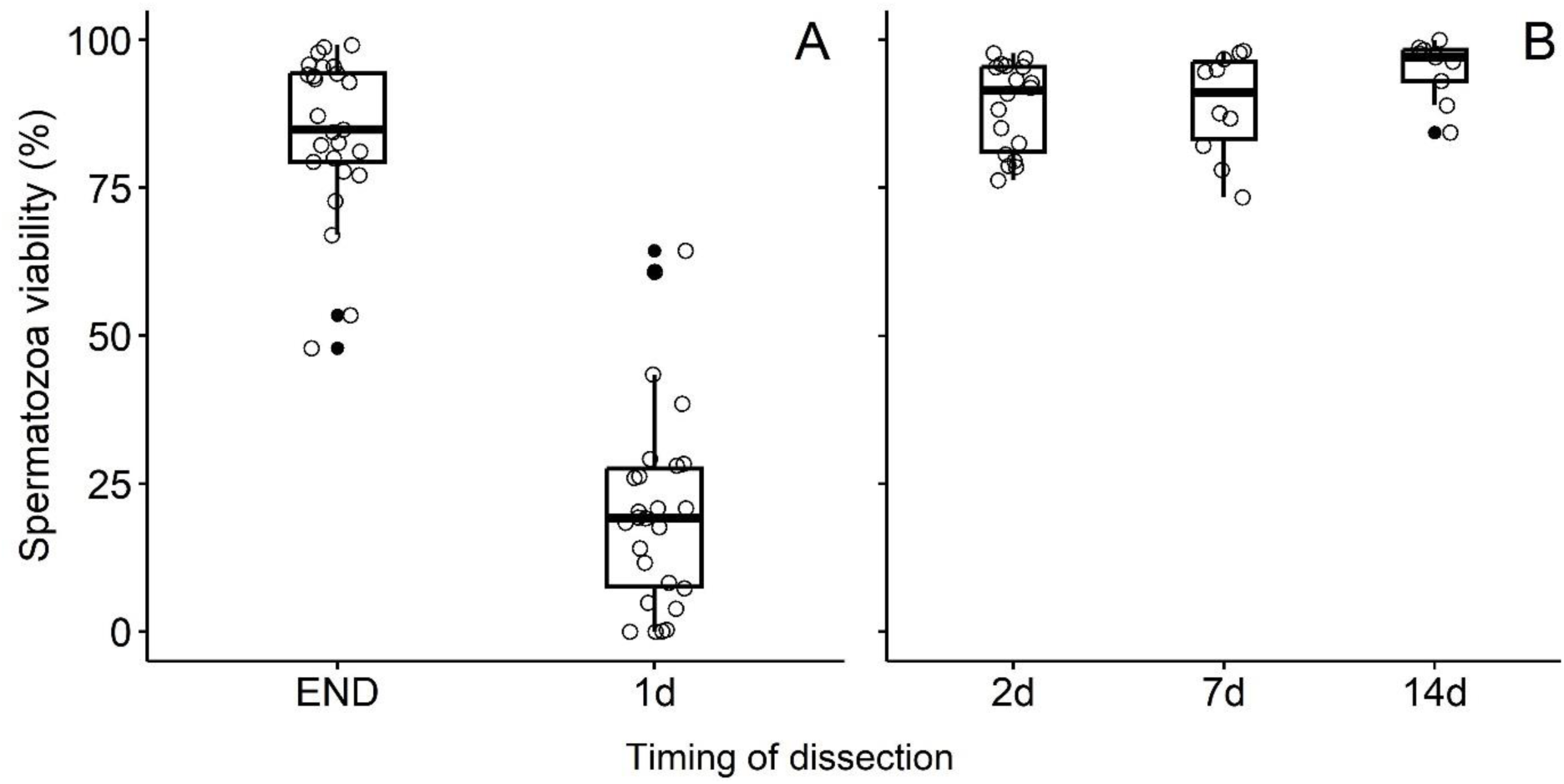
Percentage of viable spermatozoa counted (A) in the fishnet canals and (B) in the reservoirs according to the timing of dissection (*END* end of mating, *1d* one day, *2d* two days, *7d* seven days and *14d* fourteen days after mating). Box plots show median (horizontal bars), upper, and lower quartiles (borders of the box). Whiskers extend from the 10th to the 90th percentiles.

### (d) Microscopy

The fishnet canals structure consists of an epithelium that surrounds a matrix within which sperm are transferred (figure 5A). The matrix is a thin sleeve floating in the epithelium when empty (Fig 5A), leaving a space between it and the epithelium. As sperm are transferred, it is swollen to the point of being in contact with the epithelium (figure 5B), the inner layer of which is covered with microvilli (figure 5F, 5J). Two days after mating, it seems that the matrix is still in contact with the microvilli (figure 5C). The outer layer of the epithelium is concentrated in mitochondria (figure 5D, 5G) and features electron-concentrated structures resembling lysosomes (figure 5G).

**Figure 5.**
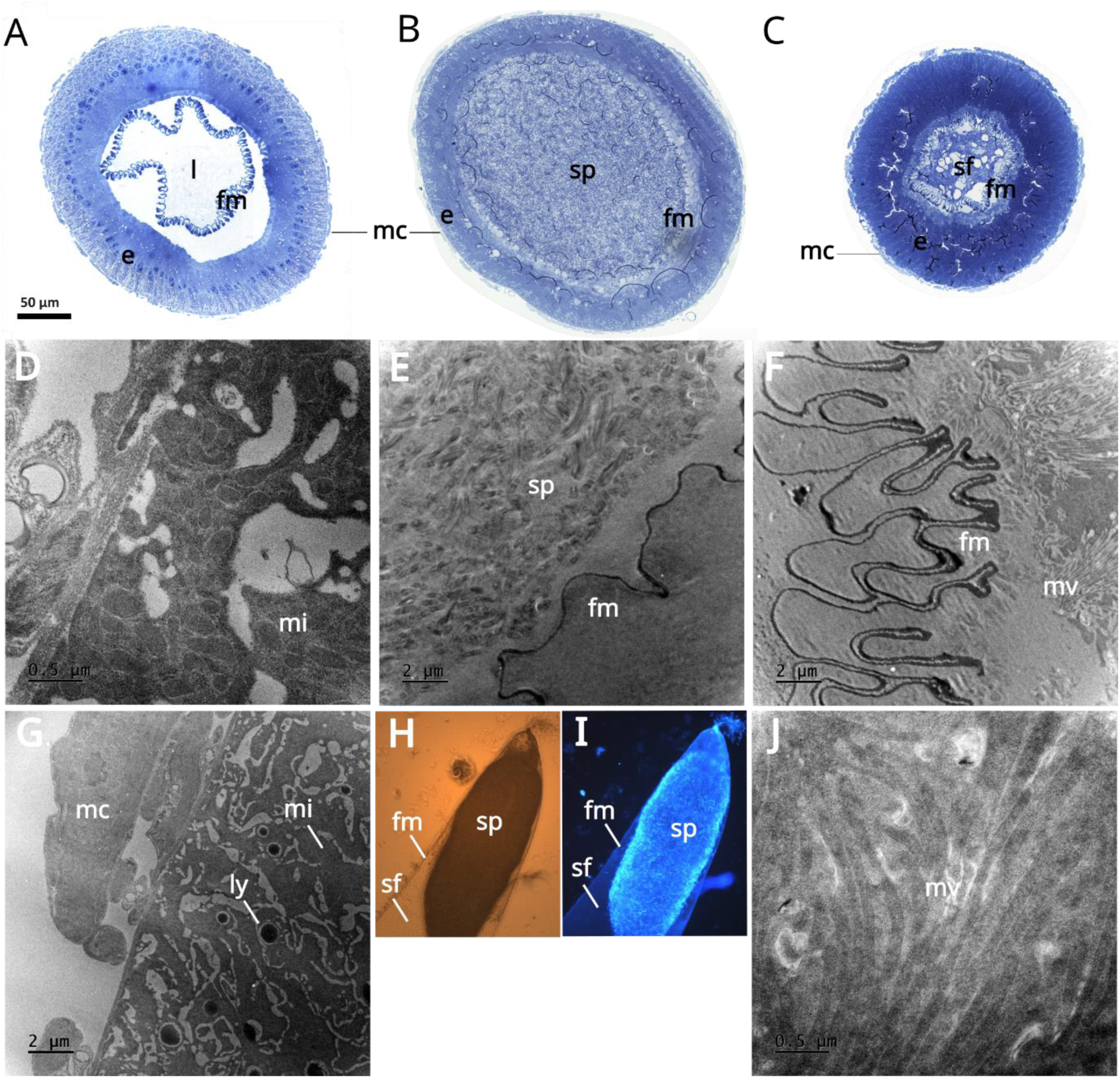
Semi-thin transversal sections of the fishnet canals (A) of a virgin female, (B) a female that just finished mating, (C) a female that mated two days before. (D) and (G): outer part of the epithelium of a fishnet canal from a virgin female. (E) The inner part of a fishnet canal from a female that just finished mating. (H) and (I): light microscopy images of the sperm plug formed at the end of the fishnet canal with visible light and fluorescence – DAPI staining was used for (I) to show spermatozoa nuclei. (F) and (J): fishnet canal matrix and microvilli of the epithelium from a female that mated two days before. Abbreviations: *fm* fishnet matrix, *e* epithelium, *l* lumen, *mc* muscular cells, *sp* spermatozoa, *sf* seminal fluid, *mi* mitochondria, *mv* microvilli, *ly* lysosomes-like structures.

Two days after mating, the fishnet canals are not fully empty as remains of seminal fluid can be seen (figure 5C). A proportion of the transferred spermatozoa are then in the reservoirs (figure 6A, 6B) where they are concentrated on one side of the capsule, leaving a space that may be filled with fluid. The reservoir wall cells are full of vacuoles (figure 6A, 6C) and structures that may have secretory properties (figure Fig. 6D).

**Figure 6.**
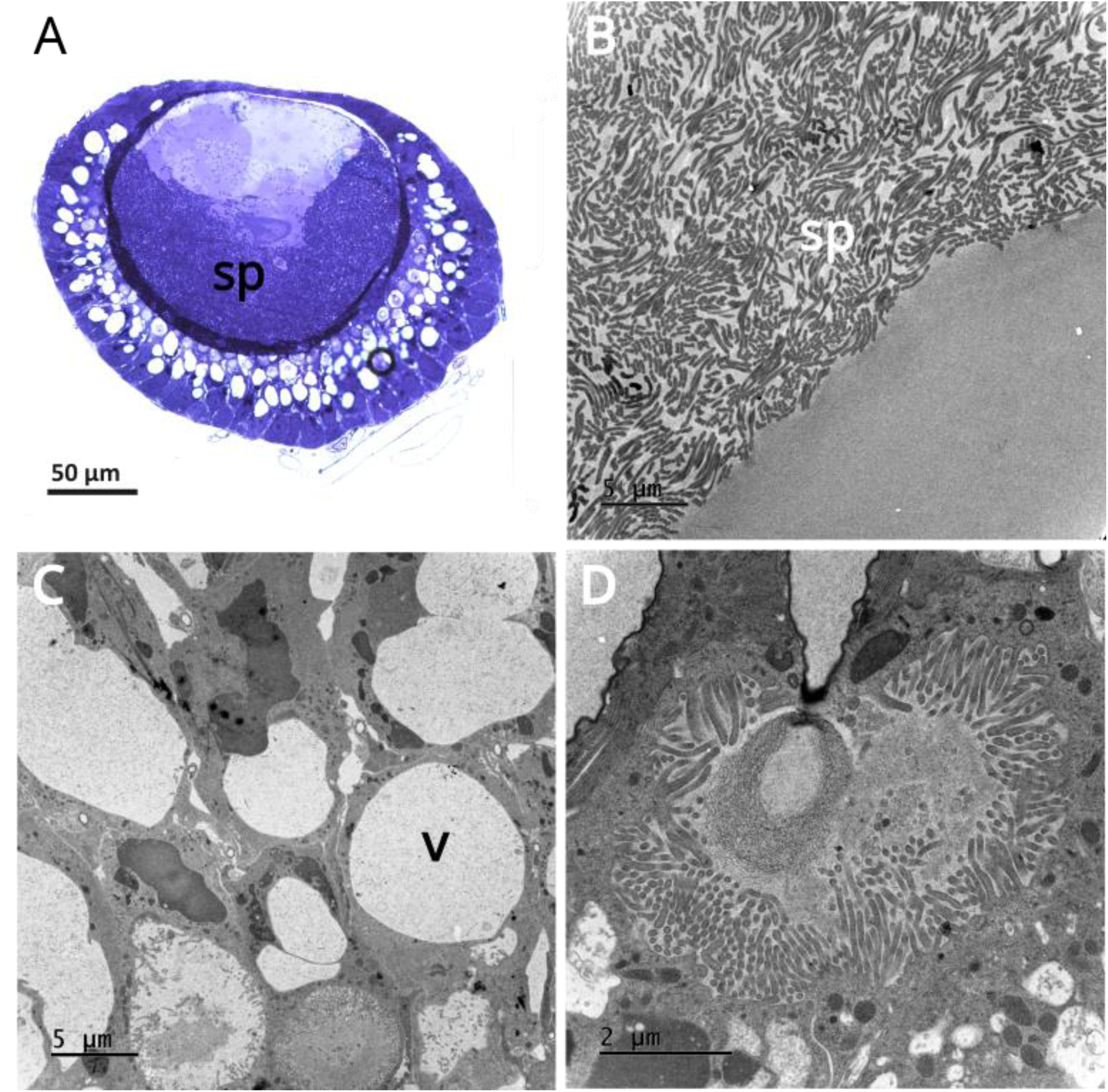
Ultrastructure of the reservoirs of a female two days after mating. (A) semi-thin section of the reservoir, (B) lumen of the reservoir full of spermatozoa, (C) ultrastucture of the reservoir capsule, (D) possible secretory compartment in the reservoir capsule. Abbreviations: *sp* spermatozoa, *v* vacuole.

## 4. Discussion

Sperm transfer in BSF appears to take place in two steps. The first one occurs during at least the first 15 minutes of mating and consists of the male transferring a seminal fluid devoid of spermatozoa which will swell the fishnet canals. The transfer of spermatozoa in the female’s spermathecae, more specifically in the fishnet canals begins after 10 minutes of mating. In insects, seminal fluids are known to have considerable impacts on female physiology – among other traits, they affect oviposition rate, sexual receptivity, sperm storage [37–39]. Here, the seminal fluids looked more gelatinous in the spermathecae than they did when being ejaculated by the male (figure S4). Some of these fluids play the role of mating plugs that retain the ejaculate within the female tract as they coagulate [11,12,40]. It seems that a similar process could take place in the spermathecae of BSF female. Whether this fluid could prevent sperm flushing or serve as a swimming support for spermatozoa is not known.

Interestingly, spermatozoa will accumulate and get stuck at the end of the fishnet canals during and after mating for one to two days before finally reaching the reservoirs of the spermathecae (contrary to what has been stated in [28]). Spermatozoa of BSF are long – 3 mm [28], and their accumulation before the access to the reservoirs resembles a mating plug made of spermatozoa - whose flagella making it look like a ball of wool (figure 5H, 5I). This apparent sperm plug may prevent the arrival of sperm from other males to the reservoirs in a context of multiple mating. It is particularly noteworthy that not all spermatozoa transferred by a male reach the reservoirs. This restricted access to the reservoirs strongly suggests that the junction between the fishnet canals and the reservoirs is a suitable site for female control over the amount of spermatozoa that go into the reservoirs.

Considering the fact that, in total, spermathecae contained more spermatozoa at the end of mating than 2 days after, it would seem that they are partly emptied between mating and oocytes fertilization, with possible sperm dumping at work here. Actually, the fishnet canals are completely emptied from spermatozoa and partially from seminal fluid one to two days after mating. However, while ejected sperm can be found in species that are known to do sperm dumping [3,33], such dumped sperm has not been observed in these experiments - either in petri dishes where females were stored or in “cages” made of microscope slides (as in [33]).

Curiously, TEM observations showed that the first part of the spermathecae, the fishnet canals, are structured like a digestive tube. The fishnet canals are composed by muscular cells surrounding an epithelium and a matrix looking like a digestive peritrophic matrix [41,42]. The epithelium showed electron-dense structures that could be lysosomes as well as microvilli— structures that are characteristics of secretory and absorbatory cells—surrounding the matrix full of sperm. Alongside the decreasing sperm number in the fishnet canals, sperm viability tests showed that the percentage of viable spermatozoa remaining in the fishnet canals decreased drastically the day after mating. Such a phenomenon has already been demonstrated in *Aedes aegypti* where it has been suggested to be the result of sperm digestion by the female [43]. The degradation of sperm observed in the fishnet canals may stem from a natural aging process of sperm in a suboptimal environment. However, taken together—the structures observed through TEM, the disappearance of sperm from the fishnet canals that has not been found to be dumped, the altered viability of spermatozoa in the fishnet canals the day after mating—, these results strongly indicate a phenomenon of sperm digestion. Given the extended lifespan observed in female BSF following mating, as documented by [29], it’s conceivable for spermatozoa, seminal fluid, or both to serve as nuptial gifts in BSF.

After mating, spermatozoa stored in the reservoirs were not a limiting resource for females fertility. Indeed, 54 % of the females laid at least two clutches among which two females laid 3 fertile clutches. All the females still had enough spermatozoa to lay at least one more clutch, and the mean number of spermatozoa in the reservoirs after one clutch would have been enough to lay approximately 4 more clutches. Moreover, sperm viability in the reservoirs remained constant over time. TEM observations of the reservoirs showed secretory cells (figure 6) whose role would certainly be to maintain spermatozoa in a suitable environment. Polyandry has been showed in this species [25,44], and even though the possibility of more clutches being laid in natural conditions can not be ruled out, sperm limitation—either by their number or their viability—does not seem to be an explanation for these multiple matings. It can be hypothesize that, if a female’s capacity to lay clutches is not constrained by the quantity or the quality of stored sperm, the potential nuptial gifts transferred by males could serve as a motivation for multiple mating [45,46].

While larger females have been found to lay more eggs—which is common in many species, including insects [47], a counterintuitive observation is the negative relationship between male size and the amount of laid eggs, even though male size is correlated with the amount of transferred seminal fluid. Furthermore, despite the positive correlation between male and female longevity, the lack of a significant relationship between male size and female longevity complicates the direct establishment of a link between the quantity of seminal fluid transferred by the male and its nutritional value.

Here, in addition to presenting the potential use of a sperm plug in BSF, a body of clues suggests that the male’s sperm could serve as an endogenous genital nutritive gift digested by the female directly in her spermathecae. The process described here represents a breeding ground for sexual conflicts [48], as strategies that are or may be used by both sexes—sperm plug in males and sperm digestion in females—may not maximise the fitness of their partner. Interestingly, if nuptial gifts are common in insects [30], endogenous genital “nutritive” gifts [49] that could be used by the female to sustain her metabolic activities are less common [49]. These findings reinforce the interest of BSF in the study of sexual selection.

## Data, scripts, code, and supplementary information availability

The data, script and supplementary materials used in this manuscript are provided by Frédéric Manas: https://doi.org/10.5281/zenodo.11402258

## Acknowledgments

We thank Hélène Girotvergne for technical assistance. We would like to thank to the IBiSA Electron Microscopy Facility of University of Tours and the University Hospital of Tours for their assistance.

